# Data paper: FoRAGE (Functional Responses from Around the Globe in all Ecosystems) database: a compilation of functional responses for consumers and parasitoids

**DOI:** 10.1101/503334

**Authors:** Stella F. Uiterwaal, Ian T. Lagerstrom, Shelby R. Lyon, John P. DeLong

## Abstract

Functional responses – the relationships between consumer foraging rate and resource (prey) density – provide key insights into consumer-resource interactions and predation mechanics while also being a major contributor to population dynamics and food web structure. We present a global database of standardized functional response parameters extracted from the published literature. We refit the functional responses with a Type II model using standardized methods and report the fitted parameters along with data on experimental conditions, consumer and resource taxonomy and type, as well as the habitat and dimensionality of the foraging interaction. The consumer and resource species covered here are taxonomically diverse, from protozoans filtering algae to wasps parasitizing moth larvae to wolves hunting moose. The FoRAGE database (doi:10.5063/F17H1GTQ) is a living data set that will be updated periodically as new functional responses are published.

## INTRODUCTION

The strength of a consumer-resource interaction determines the importance of a given food web link (Novak and Wootton, 2010). Thus, interaction strength can give key insights into population dynamics and the structure and stability of food webs (Gilbert et al., 2014; McCann et al., 1998). One way to measure consumer-resource interaction strength is through the functional response (Holling, 1959a). A consumer’s functional response describes foraging rate as a function of resource availability. The simplest form is Type I, where foraging increases linearly with resource density. Most consumers, however, must pay a time cost for each resource item utilized, resulting in an asymptotic, Type II functional response. A common form of Type II functional response is the Holling disc equation:

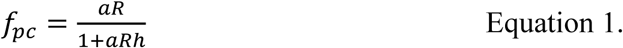

where *f*_*pc*_ is the per capita foraging rate of the consumer (number of resources per time per predator), *a* is the space clearance rate (space per time per predator), *R* is the initial resource density (resources per space), and *h* is handling time (time per resource) (Holling, 1959a). Space clearance rate describe how quickly a consumer can remove resources from a given space, while handling time describes the loss in search time associated with the consumption of an individual resource item. Handling time reflects any actcivity that prevents the consumer from searching for additional resources (prey items) after catching a resource. This may include transporting the resource to a safe location, chewing the resource, and any time spent digesting (if the consumer cannot continue hunting while digesting).

Here, we present standardized functional response parameters and the associated experimental conditions for over 2,000 consumer-resource combinations from the literature. The database will be periodically revised and updated with new functional responses as they are published. The data set is housed at the Knowledge Network for Biocomplexity (https://knb.ecoinformatics.org/) and can be found through its DOI: doi:10.5063/F17H1GTQ. The data set contains two files, one containing the overall data set of functional response parameters and associated information (with ‘data set’ in the file name) and the other containing the raw foraging observations (with ‘original curves’ in the file name).

## METHODS

We searched the literature using terms such as “functional response”, “predator-prey interaction”, and “biocontrol” to find papers reporting predator and parasitoid functional responses. We also searched within references of papers that contained functional responses, through other compilations of functional responses (DeLong et al., 2015; DeLong and Vasseur, 2012a, 2012b, 2011; Kalinoski and DeLong, 2016; Rall et al., 2012; Uiterwaal and DeLong, 2018), and through the websites of researchers that had done multiple functional response papers. Our search produced 2,083 functional responses across a wide range of taxa from all habitats and biomes and from around the world. See Appendix A for sources. Each individual functional response curve received a unique ID number associated with its functional response data, consumer and resource traits, and experimental conditions. Multiple distinct curves – due to experimental treatments, age and/or sex, consumer and/or resource species, temperature, number of consumers, substrate, or other category – from the same paper received different IDs.

### Functional response data

#### Data collection

Due to discrepancies in terms of equations and techniques used to calculate functional responses across studies, species, and experimental approaches, we did not use the parameters reported in the original papers. Rather, we recorded original resource density and consumer foraging rate data as given in the paper. We preferentially recorded raw data in tables, but we digitized most data from figures. When raw data was not available, we recorded mean foraging rate at a given resource density along with standard errors and sample size at that density. We converted error bars presented as 95% confidence intervals or standard deviation to standard error. In some cases where raw data was reported in figures, it was unclear how many actual observations were represented by a point on the graph. In these cases, we used the minimum possible number of replicates based on the reported sample size to obtain a conservative number of datapoints. If resource density was expressed in terms of carbon, protein, or chlorophyll concentration, we converted this to number of individuals using conversions provided in the paper (preferentially) or from outside sources. The data collection process is outlined in Figure 1. Additional functional response data can be included in the FoRAGE data set by making the corresponding author aware of a new source via email.

**Figure 1.**
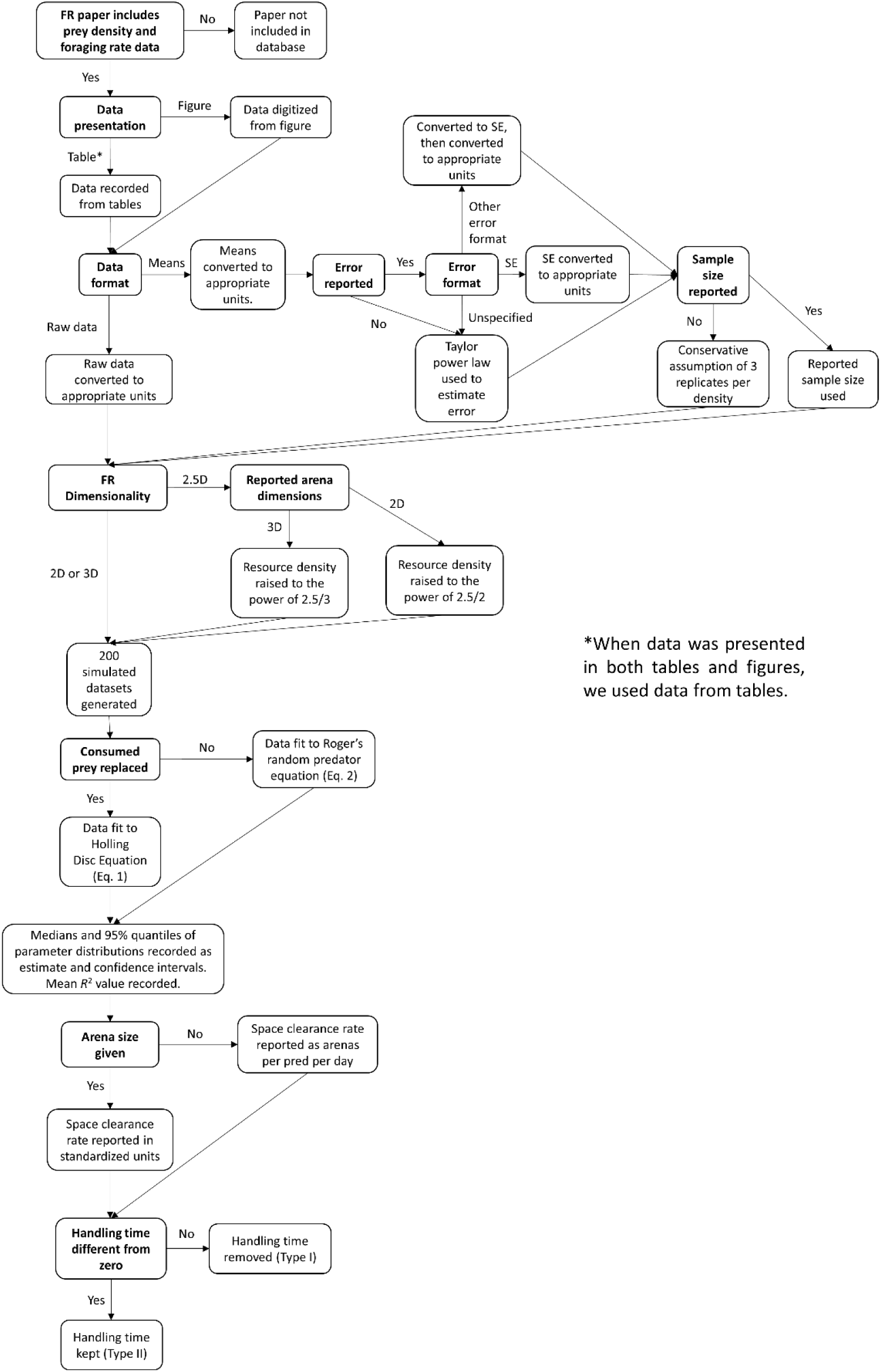
Flow diagram detailing the decisions for how to include and standardize functional response data from the literature.

#### Data fitting

We made all data comparable by standardizing resource density to units of resources per cm^2^ or m^2^ or resources per cm^3^ or m^3^ and foraging rate to units of number of resources eaten (or parasitized) per consumer per day. For 2.5-dimensional foragers (See “Experimental conditions” below), we rescaled prey density by raising it to the power of 2.5/2 if density was reported as an area and to the power of 2.5/3 if density was reported as a volume. In this way, the units of space clearance rate for organisms foraging in a fractal dimension between a plane and a volume are m^2.5^ per predator per day. However, the prey density data are retained in either two or three dimensional units within the ‘original curves’ file.

We generated 200 bootstrapped datasets per functional response. We used standard ransom sampling with replacement for data sets with raw data. For data sets that reported mean foraging rate at different densities, we generated simulated data sets with the same mean, standard error, and sample size as the reported data set using the same bootstrapping procedure. Then, we fit foraging data to the Holling disc equation (Equation 1) if the resource was replenished by the experimenters as it was consumed throughout the experiment. Datasets from wild consumers were treated as having resources replenished. If resources were not replenished, we fit foraging data to the Roger’s random predator equation:

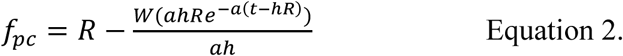

where *f*_*pc*_ is the per capita number of resources eaten by the consumer in the time of the foraging trial *t, W* is the Lambert W function, and *R, a*, and *h* are as in Equation 1. This equation accounts for resource depletion as the consumer forages (Bolker, 2008; Rogers, 1972). We used nonlinear ordinary least squares regression in Matlab to conduct these fits. To obtain parameter estimates and confidence intervals, we used the medians and 95% quantiles of the bootstrapped parameter distributions, respectively. We did not account for prey or predator population growth in the fitting procedure (Rosenbaum and Rall, 2018), although most experiments were conducted for periods precluding changes in prey or predator through reproduction.

When sample size was not given for data presented as means, we assumed conservatively that there were three replicates, the minimum number required to obtain a standard error. To estimate error when none was given or when error type was not specified, we used a Taylor power law relationship between the mean and the foraging rate variance across all observations in the data set for which this was available. We estimated this relationship using ordinary least squares regression with the log of the standard error as the dependent variable and the log of the mean foraging rate as the independent variable (Figure 2). The fitted exponent was 0.83 (0.006 ± SE) and the intercept was -1.76 (± 0.028). When arena size was not given, we conducted the fits and reported the handling time in the standard units but space clearance rate with units of ‘arenas per predator per day’. If handling time was not different from zero (95% confidence intervals overlapped zero), the functional response was assumed to be Type I and handling time was removed.

**Figure 2.**
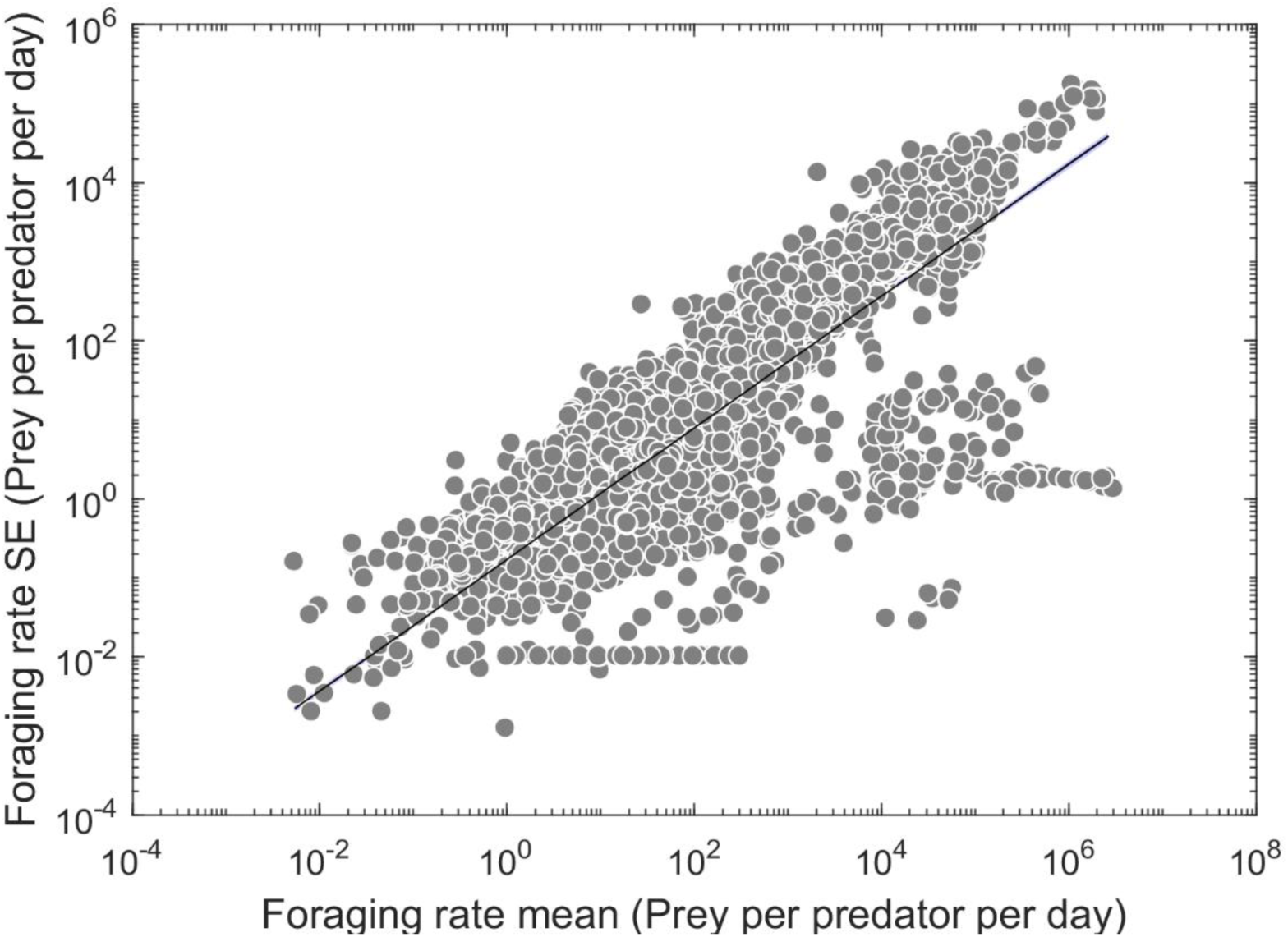
Taylor power law relationship between mean foraging rate and the standard error of the mean across all observations. This relationship was used to estimate the foraging rate standard error for studies where only the mean foraging rate at a resource level was given.

### Consumer and resource traits

#### Mass

We recorded masses or lengths of consumer and resource species from the original papers if available. If size measurements were not available in the original paper, we used mass or length estimates from external sources. We converted lengths to masses using length-weight relationships (usually by order or family). When no species-level size estimates were available, we calculated size based on one or more closely related species (e.g., in the same genus). In some cases, we were unable to obtain sizes of juveniles. In these cases, we calculated juvenile:adult size ratios of a related organism for which we did have juvenile size data. We used this percentage to estimate the unknown juvenile size of the focal organism. When necessary, we used water content to calculate wet mass from dry mass. Because predators and parasitoids utilize live resources, wet mass provides a more accurate size estimate than dry mass for functional response experiments. We used volume estimates and the density of freshwater or saltwater to estimate masses of some aquatic organisms (e.g. single-celled algae). Our methods for recording consumer and resource masses are outlined in Figure 3.

**Figure 3.**
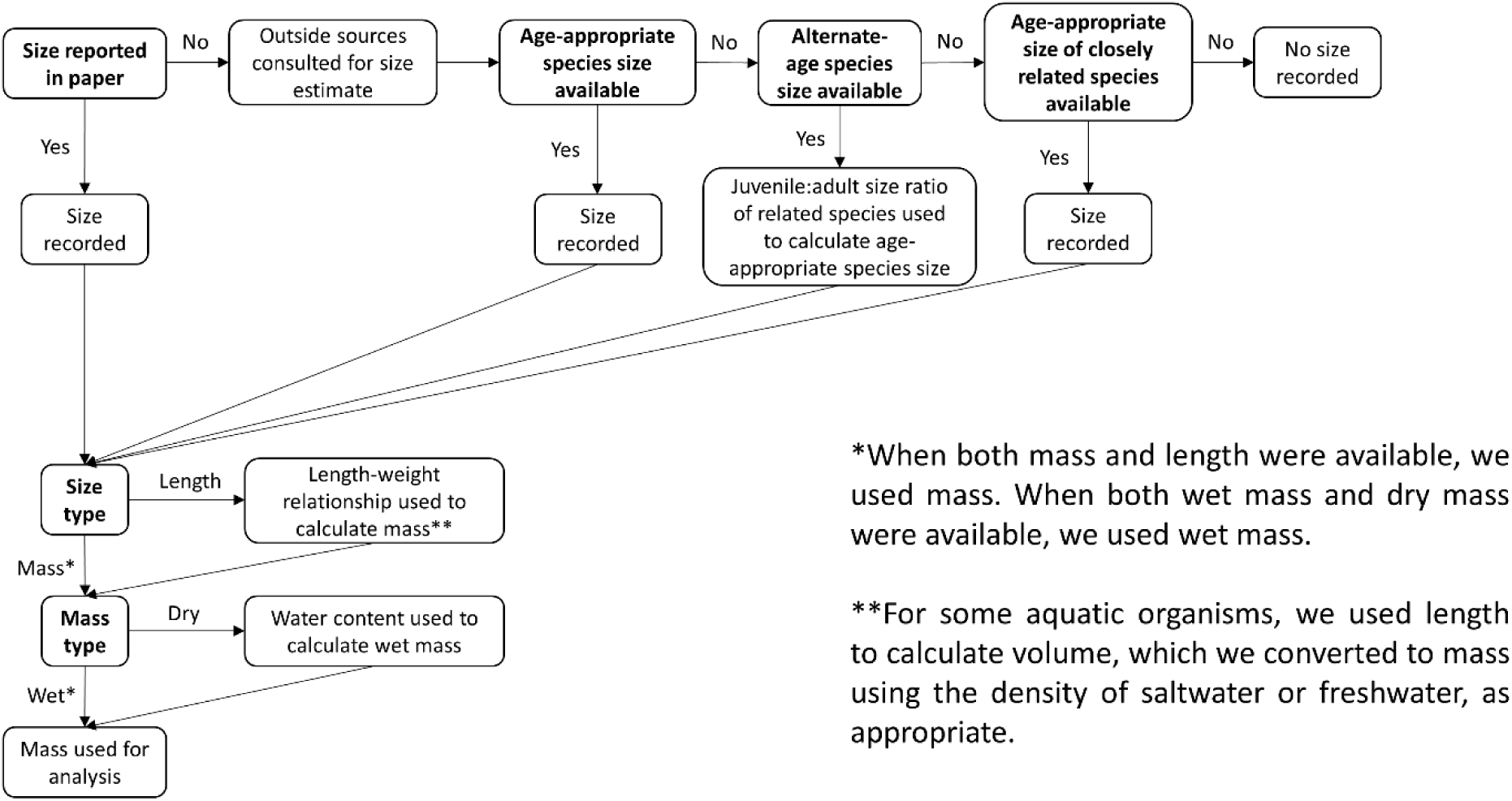
Flow diagram detailing the methods for obtaining body masses of consumers and resources represented in functional response data in the FoRAGE database.

**Figure 4.**
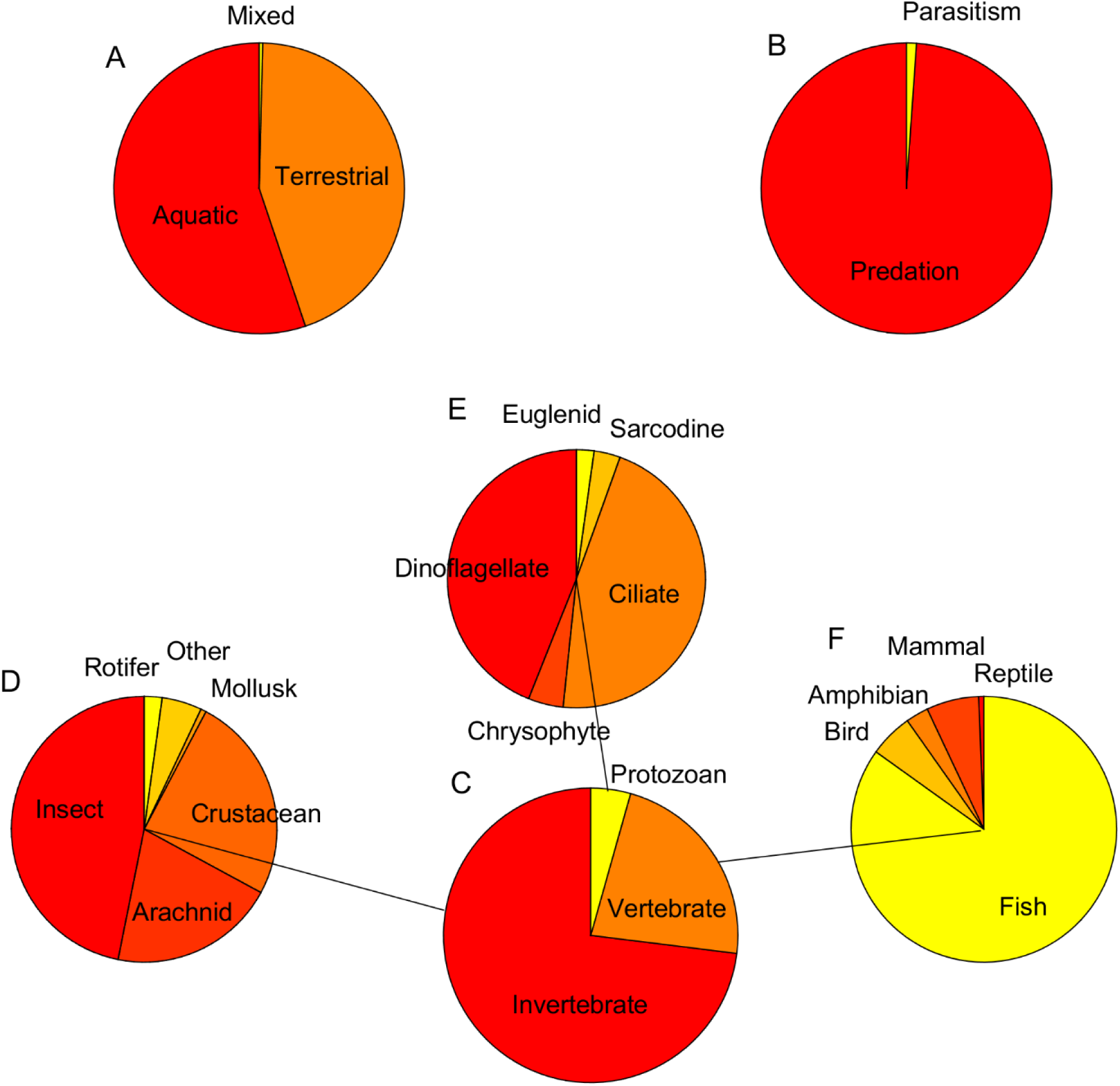
Pie charts showing the breakdown of functional responses in the FoRAGE database by A) habitat, B) foraging type, C) predator major grouping, and minor groupings within the major groupings, connected by lines, for D) invertebrates, E) protozoa, and F) vertebrates.

#### Other traits

We recorded consumer and resource traits as reported in the paper, including developmental stage, sex, and interaction type (predation or parasitoidism). We also assigned each consumer and resource to approximate taxonomic categories, including their level of cellularity (unicell or metazoan), vertebrate status (vertebrate, invertebrate, or protozoan), and other levels that represented major groupings of data such as ‘fish’ or ‘copepod’ or ‘dinoflagellate’.

#### Experimental conditions

We recorded temperature or average temperature (if a range was given) at which experiments with ectotherms were conducted. If the consumer was an endotherm, we recorded body temperature. We also reported the number of consumers per arena and the length of time consumers were deprived of resources before the experiment. We recorded notes on other experimental variables such as habitat complexity or chemical pretreatment of organisms. We also identified field-determined functional responses as ones where consumers were foraging in natural field settings rather than in a laboratory.

We determined whether the functional response occurred in two-dimensional (e.g. wolf spiders in a petri dish), three-dimensional (e.g. copepods and *Daphnia* in a tank), or 2.5-dimensional (e.g. insects crawling on whole plants, spiders on webs) space. In cases where the consumer moved in three dimensions but the resource moved in two dimensions (e.g. fish consuming bottom-dwelling crayfish), we determined the interaction to be occurring in two dimensions.

## ACKNOWLEDGEMENTS

This work was supported by a James S. McDonnell Foundation Complex Systems Scholar Award to J.P.D., a Binational Science Foundation (BSF) grant (number 2014295), and a National Science Foundation Graduate Research Fellowship to S.F.U (DGE-1610400).

---

**Appendix A:**
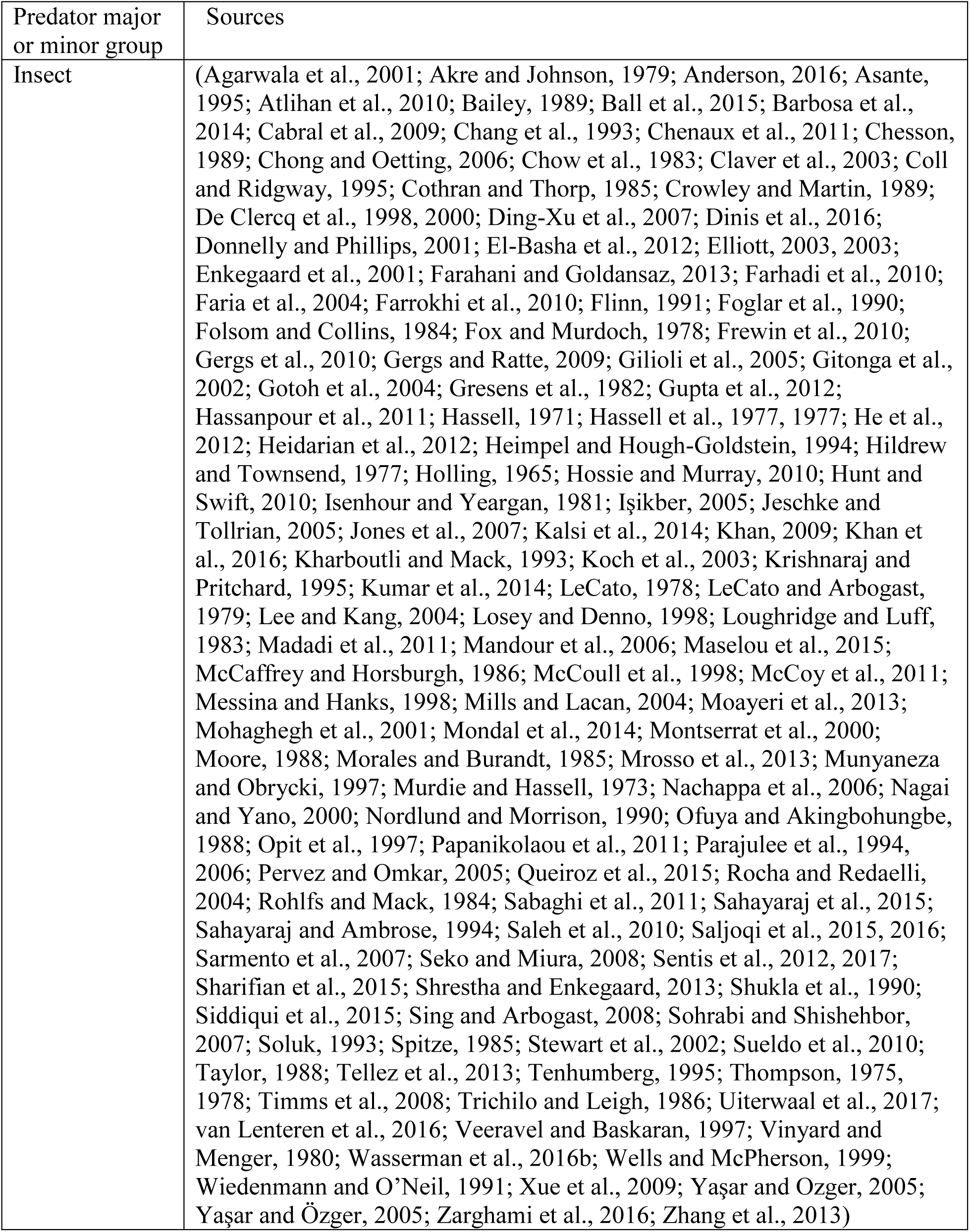
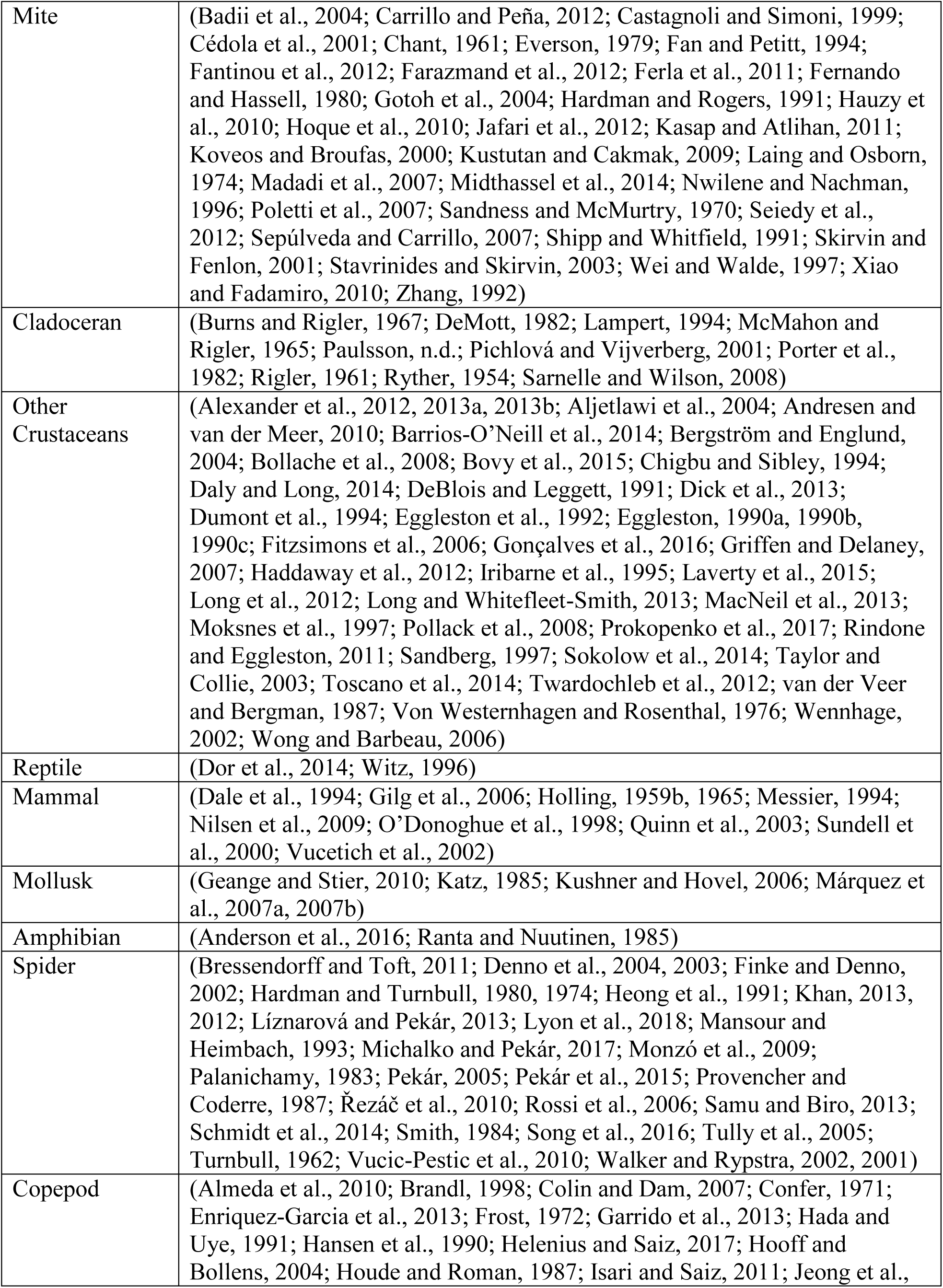
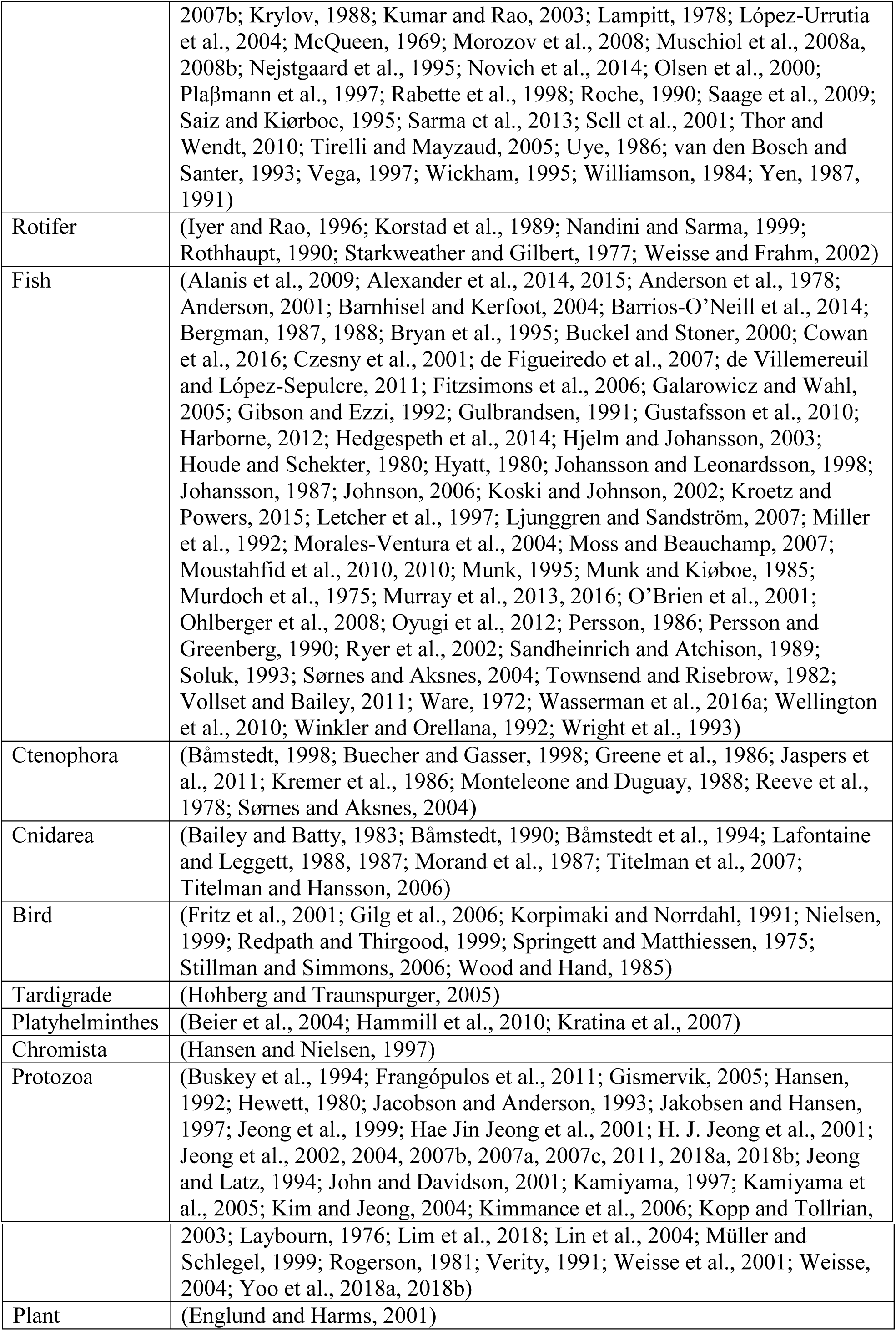
Publications containing original data on the functional responses in the FoRAGE database. Sources are grouped by Predator Major or Minor group.

